# DNA replication and chromosome positioning throughout the interphase in three dimensional space of plant nuclei

**DOI:** 10.1101/2020.04.02.021857

**Authors:** Němečková Alžběta, Veronika Koláčková, Vrána Jan, Doležel Jaroslav, Hřibová Eva

## Abstract

Despite the recent progress, our understanding of the principles of plant genome organization and its dynamics in three-dimensional space of interphase nuclei remains limited. In this study, DNA replication timing and interphase chromosome positioning was analyzed in seven *Poaceae* species differing in genome size. A multidisciplinary approach combining newly replicated DNA labelling by EdU, nuclei sorting by flow cytometry, three-dimensional immuno-FISH, and confocal microscopy revealed similar replication timing order for telomeres and centromeres as well as for euchromatin and heterochromatin in all seven species. The Rabl configuration of chromosomes that lay parallel to each other and their centromeres and telomeres are localized at opposite nuclear poles, was observed in wheat, oat, rye and barley with large genomes, as well as in *Brachypodium* with a small genome. On the other hand, chromosomes of rice with a small genome and maize with relatively large genome did not assume proper Rabl configuration. In all species, the interphase chromosome positioning inferred from the location of centromeres and telomeres was stable throughout the interphase. These observations extend earlier studies indicating a more complex relation between genome size and interphase chromosome positioning, which is controlled by factors currently not known.

**Highlight:** Telomere and centromere replication timing and interphase chromosome positioning in seven grass species differing in genome size indicates a more complex relation between genome size and the chromosome positioning.

## Introduction

One of the exciting features of eukaryotic genomes is their organization in three-dimensional space of cell nuclei and changes during cell cycle. The chromatin organization and positioning of individual chromosomes in the nucleus was a great enigma until recently. Nevertheless, based on microscopic observations of dividing cells of *Salamandra maculata* and *Proteus anguinus*, Carl Rabl predicted already in 1885 that the positioning of chromosomes in interphase nuclei follows their orientation in the preceding mitosis (Reviewed by Cremer *et al.*, 2006). The hypothesis was confirmed later by cytogenetic studies in human, animals as well as in plants with larger chromosomes, which demonstrated that centromeric and telomeric regions cluster at opposite poles of interphase nuclei (Cremer *et al.*, 1982; Schwarzacher and Heslop-Harrison 1991; Werner *et al.*, 1992). This arrangement of chromosomes in interphase was dubbed the Rabl configuration.

Interestingly, a different arrangement of chromosomes in interphase nuclei was observed in *Arabidopsis thaliana*, a model plant with small genome (1C~157 Mbp). Here, the centromeres are located at the nuclear periphery, whereas the telomeres congregate around nucleolus (Armstrong *et al.*, 2001; Fransz *et al.*, 2002). Centromeric heterochromatin forms dense bodies called chromocenters, while euchromatin domains form 0.2 - 2 Mb loops that are organized into Rosette-like structures (Fransz *et al.*, 2002; Pecinka *et al.*, 2004; Tiang *et al.*, 2012; Schubert *et al.*, 2014).

Some earlier studies suggested that chromatin arrangement in interphase nuclei of wheat, oat and rice, and in particular the centromere - telomere orientation, may be tissue-specific and cell cycle-dependent (Dong and Jiang 1998; Prieto *et al.*, 2004; Santos and Show, 2004). While a majority of nuclei in somatic cells of rice (*Oryza sativa*), which has a small genome (1C~490 Mbp, Bennett *et al.*, 1976) lack Rabl configuration, chromosomes in the nuclei of some rice tissues, e.g., pre-meiotic cells in anthers or xylem - vessel precursor cells seem to assume the configuration (Prieto *et al.*, 2004; Santos and Show 2004). Such chromosome arrangement and orientation in somatic cell nuclei was also observed in other plants with small genomes, including *Brachypodium distachyon* (1C~ 355 Mbp) (Idziak *et al.*, 2015). Although the majority of cell types of small plant genomes of *B. stacei* (1C~276 Mbp, Catalán *et al.*, 2012) and *B. hybridum* (1C~619 Mbp, Catalán *et al.*, 2012) did not show Rabl configuration. On the other hand, recent study of Idziak *et al.* (2015) showed that Rabl configuration is present in small proportion (13 and 17%) of root meristematic cells of both these plant species. Rabl configuration was not observed also in plants with large genomes such as *Allium cepa* (~ 1.5 Gb), *Vicia faba* (~ 15Gb), *Solanum tuberosum* (~ 900 Mbp) and *Pisum sativum* (~ 3.9 - 4.4 Gb) (Rawlins *et al.*, 1991; Fussell, 1992; Kamm *et al.*, 1995; Harrison and Heslop-Harrison, 1995).

Development of a three-dimensional fluorescence *in situ* hybridization (3D-FISH) method provided an opportunity to map telomere positions at meiotic prophase in maize (Bass *et al.*, 1997), Arabidopsis and oat (Howe *et al.*, 2013). In order to characterize changes in chromosome positioning in 3D nuclear space throughout the interphase, cell nuclei at particular cell cycle phase can be isolated using flow cytometry. Embedding flow-sorted nuclei in polyacrylamide gel stabilizes their structure during the 3D-FISH procedure and 3D images can be captured by confocal microscopy (Kotogány *et al.*, 2010; Hayashi *et al.*, 2013; Bass *et al.*, 2014; Koláčková *et al.*, 2019).

Chromosome conformation capture (3C) technique (Dekker 2002) and its variants offer an alternative approach to study spatial organization of chromatin in cell nuclei. So called Hi-C method (Lieberman-Aiden *et al.*, 2009) analyses contacts between DNA loci across the whole genome. The contact maps thus obtained enable the analysis of chromosome contact patterns, genome packing and 3D chromatin architecture (Dong *et al.*, 2018; Kempfer and Pombo, 2019). It should be noted, however, that Hi-C identifies genome loci that are associated in 3D space, but it does not provide the information on their physical position in the nuclei.

In a majority of cases, spatial organization of chromatin in interphase nuclei revealed by FISH corresponded to the results obtained by Hi-C (Sexton *et al.*, 2012; Dong *et al.*, 2017; Mascher *et al.*, 2017; Liu *et al.*, 2017). The only exception was *Arabidopsis thaliana* where Hi-C analysis did not confirm the Rosette-like organization of chromosomal domains. According to Hi-C, telomeres of different chromosomes should cluster, but FISH studies show telomeres located around nucleolus (Feng *et al.*, 2014; Liu and Weigel *et al.*, 2015).

Several investigations focused on organization of chromatin, its structure and changes during cell cycle. In mammals, DNA sequences located at interior regions of nuclei are replicated earlier, while late replication occurred mostly in nuclear periphery (Gilbert *et al.*, 2010; Bryant and Aves 2011). Cell cycle kinetics and the progression of cells through S phase can be followed in detail after labelling newly synthesized DNA by a thymidine analogue 5-ethynyl-2’-deoxyuridine (EdU) and subsequent bivariate flow cytometric analysis of nuclear DNA content and the amount of incorporated EdU (Mickelson-Young *et al.*, 2016). EdU-labelled nuclei can be sorted by flow cytometry and used as templates for FISH to analyze replication timing of particular DNA sequences and their positioning in 3D nuclear space (Hayashi *et al.*, 2013; Bass *et al.*, 2014; Bass *et al.*, 2015; Dvořáčková *et al.*, 2018).

Using 3D microscopy to analyze DNA replication dynamics in maize root meristem cells, Bass *et al.* (2015) observed distinct patterns of EdU signal distribution during early and middle S phase. In early S phase, DNA replication primarily occurred in regions characterized by weak DAPI fluorescence, while replication in middle S phase correlate with strong DAPI signals. Authors also revealed that knob regions and centromeric regions associated with heterochromatin were replicated during late S phase. Based on their findings they proposed “mini-domain model”, when gene islands are replicated in early S phase, and blocks of repetitive DNA in middle S phase (Bass *et al.*, 2015).

DNA replication dynamics was analyzed in detail by microscopy in several plant species, including monocot species maize and barley, and dicots Arabidopsis and field bean (Jasencakova *et al.*, 2001; Bass *et al.*, 2014; Jacob *et al.*, 2014; Robledillo *et al.*, 2018). In general, different stages of S phase contrasted in DNA replication pattern. The early S phase was characterized by weak dispersed signals of EdU, speckled signals of EdU were typical for late S phase, while the nuclei in the middle S phase were all covered with EdU signals, except of nucleolar area (Jasencakova *et al.*, 2001; Kotogány *et al.*, 2010; Bass *et al.*, 2014; Bass *et al.*, 2015;).

Interestingly, the dynamics of interphase chromosome positioning during cell cycle in plants was not studied to date. In order to fill this gap, we characterized chromosome positioning in 3D nuclear space during cell cycle and identified replication timing of DNA in centromeric and telomeric regions. To do this, we labelled newly synthesized DNA by EdU, followed cell cycle kinetics by flow cytometry and performed 3D immuno-FISH on flow-sorted nuclei. Chromosome positioning was analyzed in interphase nuclei of root meristem cells in seven *Poaceae* species differing in nuclear genome size. Confocal microscopy of nuclei embedded in polyacrylamide gel allowed us to locate centromeres and telomeres in 3D space and to analyze replication timing of these chromosome regions. The results provided new information on chromosome positioning and spatio-temporal pattern of DNA replication of the important chromosome domains.

## Material and methods

### Plant material and seeds germination

Plants used in the present study included wheat *(Triticum aestivum* L.) cultivar Chinese Spring (2n=2x=42), oat (*Avena sativa* L.) cultivar Atego (2x=2x=42), barley (*Hordeum vulgare* L.) cultivar Morex (2n=2x=14), rye (*Secale cereale* L.) cultivar Dánkowskie Diament (2n=2x=14), rice (*Oryza sativa* L.) cultivar Nipponbare (2n=2x=24), maize (*Zea mays* L.) line B73 (2n=2x=20) and *Brachypodium distachyon* L. cultivar Bd21 (2n=2x=10). All seeds were obtained from IPK Genebank (Gaterleben, Germany) except for rice, which was kindly provided by prof. Takashi Ryu Endo (Kyoto University, Kyoto, Japan) and wheat, which was obtained from Wheat Genetics & Genomic Resources Center (Kansas state university, Manhattan, KS 66506). All seeds were germinated in a biological incubator at 24 °C in glass Petri dishes on moistened filter paper until the primary roots were 2.5 – 4 cm long.

### EdU labelling of replicating DNA

Young seedlings were incubated in 20 μM 5-ethynyl-2’-deoxyuridine (EdU) (Click-iT™ EdU Alexa Fluor™ 488 Flow Cytometry Assay Kit, ThermoFisher Scientific/Invitrogen, Waltham, Massachusetts, USA) made in ddH_2_O for 30 min at 24 °C. Root tips were excised and fixed in 2% (v/v) formaldehyde in Tris buffer for 20 min at 4°C, and washed 3 times in Tris buffer at 4°C (Doležel *et al.*, 1992). About ten root tips with incorporated EdU were treated in 0.5 ml Click-iT reaction cocktail (Click-iT™ EdU Alexa Fluor™ 488 Flow Cytometry Assay Kit) prepared according to the manufacturer instructions, incubated for 10 min under vacuum, followed by incubation in the dark for 45 min at room temperature (RT). After the labelling, roots were washed 3 times for 5 min in phosphate buffered saline (PBS; 10 mM Na_2_HPO_4_, 2mM KH PO, 137mM NaCl, 2.7mM KCl, pH 7.4) on a rotating shaker (160 rpm) at RT.

The root tips were mounted in 0.1 % agarose made in re-distilled water onto cavity microscopic slides (Paul Marienfeld GmbH & Co. KG, Lauda-Königshofen, Germany) and mounted in Vectashield with DAPI (Vector Laboratories, Ltd., Peterborough, UK) to counterstain the chromosomes. The preparations were imaged using a Leica TCS SP8 STED 3X confocal microscope (Leica Microsystems, Wetzlar, Germany) with 10x/0.4 NA Plan-Apochromat objective (*z-stacks*, pinhole Airy). Image stacks were captured separately for DAPI using 405 nm laser and for EdU labelled by Alexa Fluor 488 using 488 nm laser, and appropriate emission filters. Typically, image stacks of 36 slides in average with 138 μm spacing were acquired. Finally, maximum intensity projections were done using Leica LAS-X software and final image processing was done using Adobe Photoshop 6 (Adobe Systems).

### EdU labelling of replicating DNA and nuclei preparation for flow cytometry

Roots of young seedlings were incubated in 20 μM EdU made in ddH_2_O (Click-iT^™^ EdU Alexa Fluor^™^ 488 Flow Cytometry Assay Kit) for 30 min at 24 °C. Suspensions of intact nuclei were prepared according to Doležel *et al.* (1992). Briefly, ~1 cm-long root tips were cut and fixed with 2% (v/v) formaldehyde in Tris buffer (10 mM Tris, 10 mM Na_2_EDTA, 100 mM NaCl, 0.1% Triton X-100, 2 % formaldehyde, pH 7.5) at 4°C and washed three times with Tris buffer at 4°C. Meristematic parts of root tips (~1 mm-long) were excised from 70 roots per sample in barley, oats, wheat, rye, and from 100 roots in rice and *Brachypodium*. Root meristems were homogenized in 500 μl LB01 buffer (Doležel *et al*., 1989) by Polytron PT 1200 homogenizer (Kinematica AG, Littua, Switzeland) for 13 s at 10 000 – 24 000 rpm depending on a species. For maize, 50 root meristems were chopped using a razor blade according to Doležel *et al*. (1989). The crude homogenates were passed through 50 μm nylon mesh and nuclei were pelleted at 500 g, at 4 °C for 10 min. The pellet was resuspended in 0.5 ml Click-iT reaction cocktail prepared according to the manufacturer instructions and the nuclei were incubated in the dark at 24 °C for 30 min. Then the nuclei were pelleted at 500 *g* for 10 min, resuspended in 500 μl of LB01 buffer and stained by DAPI (0.2 μg/ ml final concentration). Finally, the suspensions were filtered through a 20 μm nylon mesh and analyzed using FASCAria II SORP flow cytometer and sorter (BD Bioscience, San Jose, USA) equipped with a UV (355 nm) and blue (488 nm) lasers. Nuclei representing different phases of cell cycle were sorted into 1x meiocyte buffer A (1x buffer A salts, 0.32M sorbitol, 1x DTT, 1x polyamines) (Bass *et al.*, 1997; Howe *et al.*, 2013).

### Nuclei mounting in polyacrylamide gel

Flow sorted nuclei were mounted in 5% polyacrylamide (PAA) gel as described by Howe *et al.* (2013) and Bass *et al.* (1997) with minor modifications. Briefly, 500 μl PAA mix containing 15 % (w/v) acrylamide/bisacrylamide (akrylamid/bisakrylamid 30% NF ROTIPHORESE (29: 1) Roche Applied Science, Penzberg, Germany), 1x Buffer A salts (10 x buffer contains 800 mM KCl, 200 mM NaCl, 150 mM PIPES, 20 mM EGTA, 5 mM EDTA, 1 M NaOH, pH 6.8), 1x polyamines (1,000 x polyamines contains 0.15 M spermine and 0.5 M spermidine), 1x dithiothreitol (1,000 x dithiothreitol contains 1.0 M DTT, 0.01 M NaOAc), 0.32 M sorbitol and 99 μl ddH2O was rapidly combined with 25 μl of freshly prepared 20% ammonium sulfate in ddH_2_O and 25 μl 20% sodium sulfate (anhydrous) in ddH_2_O. One volume of activated PAA gel mix was mixed with two volumes of flow-sorted nuclei on a microscopic slide coated with aminoalkylsilane (Sigma-Aldrich, Darmstadt, Germany) and stirred gently with the pipette tip. The PAA drop was covered with a clean glass coverslip and let to polymerize at 37°C for ~40 min. The coverslip was removed and silane-coated slides with acrylamide pad were washed three times with 1x MBA in Coplin jar to remove unpolymerized acrylamide.

### Immuno-staining and fluorescence *in situ* hybridization (FISH)

In order to visualize centromeric regions, slides with PAA pad were washed in blocking buffer (phosphate buffer, 1% Triton X-100, 1 mM EDTA) at RT for 1 h, after which 100 μl of blocking buffer was added per slide and covered with parafilm for 10 min. Next, 50 μl of diluted anti-OsCenH3 primary antibody (1:100) (Nagaki *et al.*, 2004) was added, covered with parafilm and incubated at 4°C in a humid chamber for 12 hrs. The slides were then washed in a wash buffer (phosphate buffer, 0.1% Tween, 1 mM EDTA) three times for 15 min and once for 10 min in 2x Saline-Sodium Citrate buffer (2x SSC) and fixed in 1% (v/v) formaldehyde in 2 x SSC for 30 min at RT. After the fixation, slides were washed 3 x 15 min in 2x SSC at RT and used for FISH.

Hybridization mix (35 μl) containing 50% formamide, 2x SSC and 400 ng of directly labelled telomere oligo-probe ([CCCTAAA]_4_) was added onto a slide and covered by a glass coverslip. The slides were denatured at 94 °C for 6 min and incubated in a humid chamber at 37 °C overnight. Post-hybridization washing steps comprised of 5 min wash in 2x SSC, stringent washes 2 x 15 min in 0.1 x SSC at 37°C, 2 x 15 min in 2x SSC at 37 °C, 2 x 15 min 2x SSC at RT, and 1 x 15 min in 4x SSC at RT. After washing, 100 μl of secondary antibody diluted 1:250 in blocking buffer (5 % bovine serum albumin with 0.1 % Tween dissolved in phosphate buffer, secondary antibody anti-Rabbit Alexa Fluor 546 (ThermoFisher Scientific/Invitrogen) was added, covered with parafilm and incubated at 37 °C for 3h. Finally, the preparation was washed in 4x SSC (3 x 15 min, RT) and in phosphate buffer (3 x 15 min, RT). PAA pad was mounted in 30 μl of Vectashield with DAPI (Vector Laboratories), covered with a glass coverslip, and sealed with nail polish.

### Confocal microscopy and image analysis

Images were acquired using Leica TCS SP8 STED 3X confocal microscope (Leica Microsystems) equipped with 63x/1.4 NA Oil Plan-Apochromat objective (*z-stacks*, pinhole Airy) equipped by Leica LAS-X software with Leica Lightning module. Image stacks were captured separately for each fluorochrome using 647 nm, 561 nm, 488 nm, and 405 nm laser lines for excitation and appropriate emission filters. Typically, image stacks of 100 slides on average with 0.2 μm spacing were acquired. 3D models of microscopic images, colocalization analysis and volume calculations were performed using Imaris 9.2 software (Bitplane, Oxford Instruments, Zurich, Switzerland). To estimate colocalization signals, Imaris software’s ‘Colocalization’ function based on Pearson’s correlation coefficient were used (Manders *et al.*, 1992). The region of interest (ROI) was individually determined for each nucleus and each channel. Importantly, setting ROI ensures, the ‘layer by layer’ correlation, so negative colocalization of green channel of EdU signals representing another layer is prevented. The volume of each nucleus and EdU signals were detected based on primary intensity of fluorophores obtained after microscopic analysis. Imaris function ‘Surface’ and ‘Spot detection’ was used for modeling the centromere - telomere arrangements. Chanel contrast was adjusted using ‘Chanel Adjustment’ and videos were created using ‘Animation function’. Between 100 and 150 nuclei were analyzed per each plant species.

### Results

As we have studied plant species differing in genome sizes and root morphology, experimental protocols had to be individually optimized. While EdU concentration and incubation times were the same for all species, microscopic analysis of root tips after EdU labelling revealed differences in the area of meristematic zones (Fig. 1). This information was useful for the excision of meristematic regions to prepare suspensions of meristem cell nuclei. Other critical step of the preparation of nuclei suspensions was the extent of mechanical homogenization, which affected the nuclei integrity and yield (Table 1). The thick maize roots were chopped in nuclei isolation buffer using a razor blade to obtain sufficient amounts of nuclei suitable for flow cytometry.

**Table 1.** Parameters for nuclei suspension preparation after EdU pulse for individual species.

| **Plant species** | **Number of roots** | **Homogenisation [rpm]** | **Time [s]** |
| --- | --- | --- | --- |
| *Brachypodium distachyon* | 100 | 10000 | 13 |
| *Oryza sativa* | 100 | 14500 | 13 |
| *Zea mays* | 50 | Chopped by razor blade | - |
| *Hordeum vulgare* | 70 | 14500 | 13 |
| *Secale cereale* | 70 | 14500 | 13 |
| *Avena sativa* | 70 | 24000 | 13 |
| *Triticum aestivum* | 70 | 20000 | 13 |

**Figure 1.**
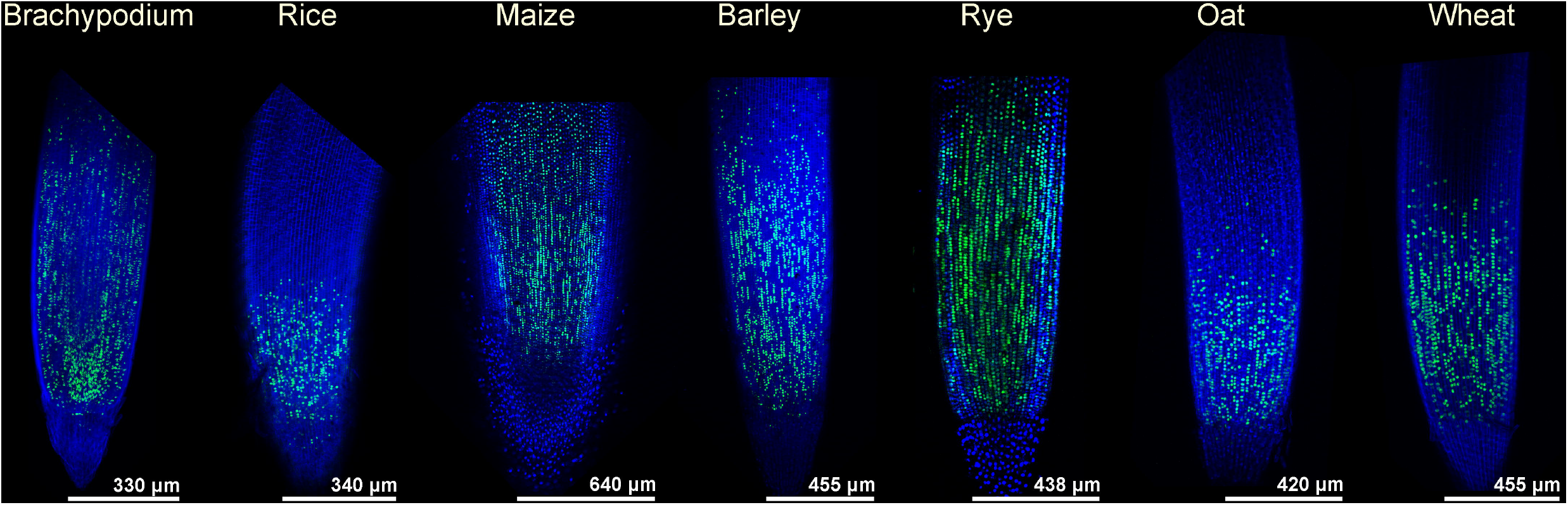
Root meristematic zones of seven *Poaceae* species. Roots were incubated with 20 mM EdU for 30 min and EdU incorporated into replicating DNA was detected by Alexa Fluor 488 (green color). Nuclei were stained with DAPI (blue color).

DNA replication kinetics in interphase nuclei of root tip meristems during cell cycle was evaluated after EdU incorporation into replicating DNA. Bivariate flow cytometric analysis EdU vs. DAPI fluorescence resulted in typical “horseshoe” dotplot patterns, which made it possible to unambiguously distinguish the nuclei at G1 and G2 phases of the cell cycle and in early, middle and late S phase (Fig. 2). Fluorescent detection of incorporated EdU was useful not only for flow cytometric analysis and nuclei sorting, but also for microscopy.

**Figure 2.**
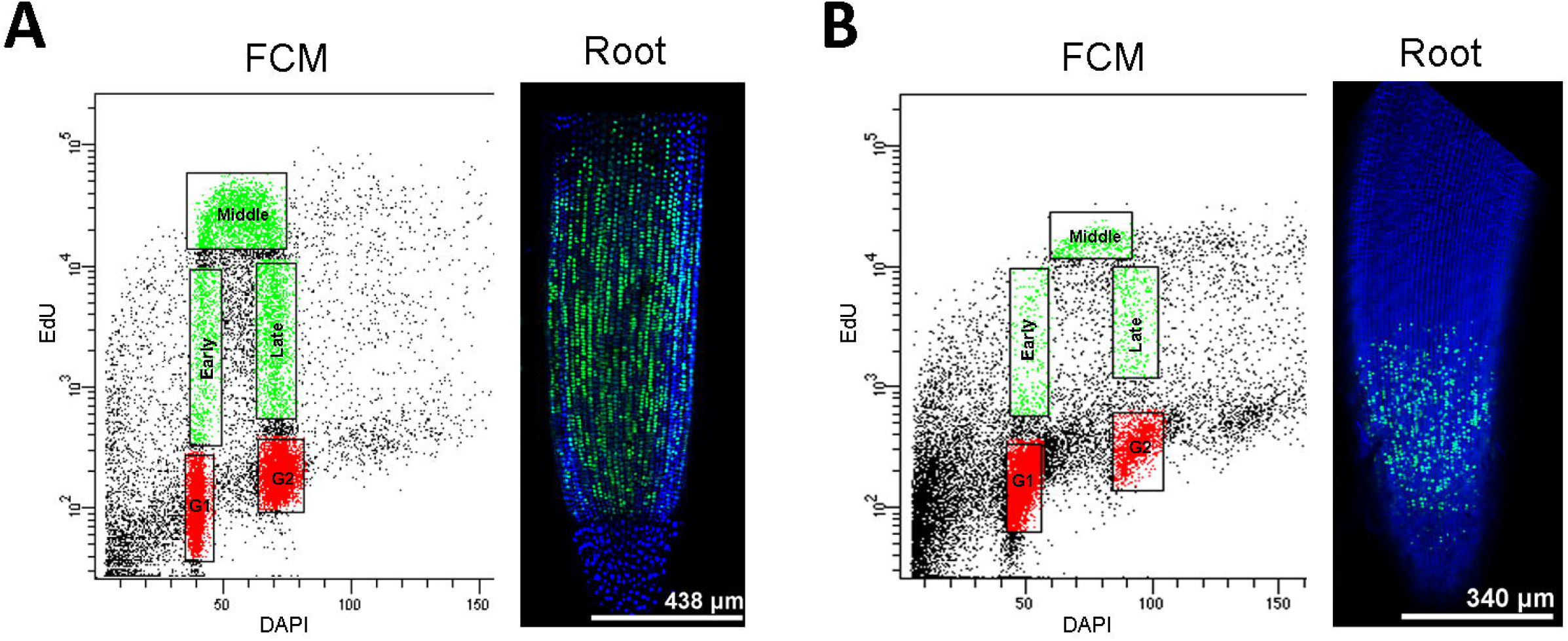
Bivariate flow cytometric analysis of cell cycle in rye (A) and rice (B). Roots of young seedlings were incubated with 20 mM EdU for 30 min. EdU in isolated nuclei was detected by Alexa Fluor 488 (green color) and their DNA was stained with DAPI (blue). ***x*** axis represents relative DNA content estimated as the intensity of DAPI fluorescence (linear scale). ***y*** axis shows the extent of EdU incorporation into newly synthesized DNA quantified by Alexa Fluor 488 fluorescence intensity (log scale). Red boxes in the dot plot show G1- and G2-phase nuclei, green boxes highlight the early, middle and late S phase.

#### DNA replication kinetics

Microscopic analysis of EdU fluorescence in cell nuclei revealed different replication patterns specific for early, middle and late S phase. Weak discrete signals were typical for early DNA replication stage, while speckled signals concentrated in particular areas were characteristic for late DNA replication stages. Strong signals dispersed throughout the whole nuclei were observed in nuclei at middle S phase (Fig. 3). Overlaps between heterochromatin regions and EdU signals at late S phase indicated that these regions were replicated later than euchromatin. EdU signals were also detected inside nucleoli as discrete and well visible spots during early and middle S phase. Clear signals were also seen at the periphery of nucleoli at middle and late S phase (Supplementary Fig. 1), suggesting the replication of 45S rDNA loci.

**Figure 3.**
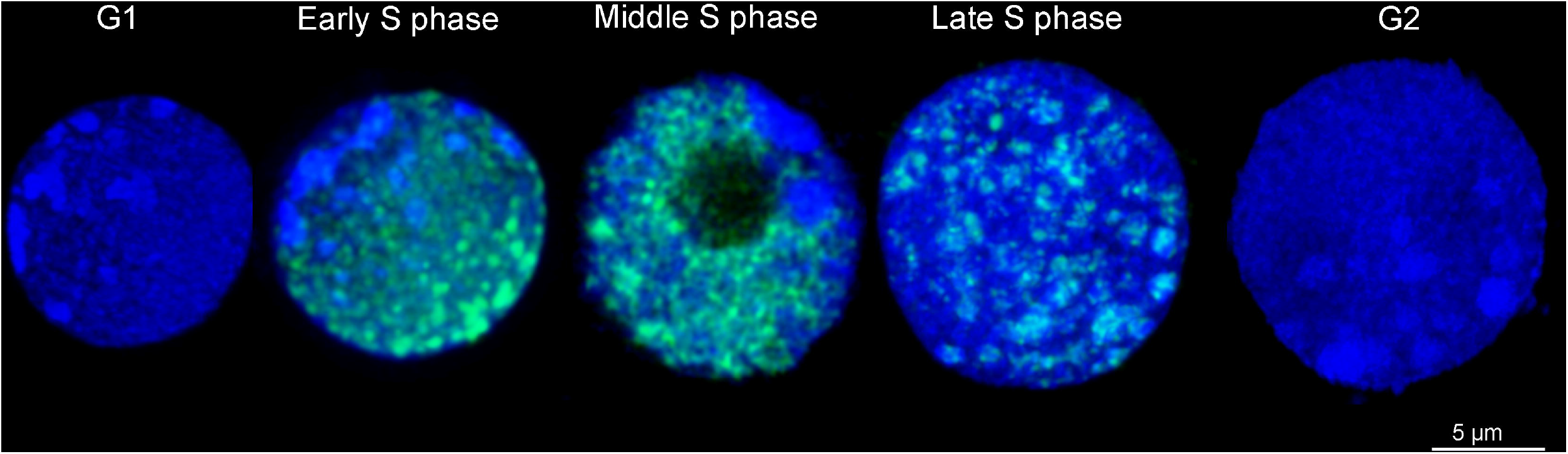
Maximum intensity projection of rye nuclei in 3D in different phases of cell cycle. EdU (green color) was incorporated during a 30 min pulse into the newly synthesized DNA. G1 and G2 phases lack green signals of EdU. Changes in DNA replication pattern during early, middle and late S phase are clearly visible. Note the replication of heterochromatin regions. DNA was stained by DAPI (blue color).

#### Replication timing of centromeric and telomeric regions

Replication timing of centromeric and telomeric regions was determined after microcopy of nuclei at different stages of S phase and based on the overlap of EdU and immunofluorescent signals at centromeric regions and FISH signals at telomeric regions (Fig. 4). Although the replication of centromeric regions initiated at early S phase, the highest number of centromeric regions underwent replication during middle and late S phase (Fig. 4, 5, Supplementary Fig. 2, 3, 4). In contrast, prevalent replication of telomeric sequences was observed during early and middle S phase. This replication pattern of both chromosome domains was observed in all species (Fig. 4, 5, Supplementary Fig. 5, 4, 7) except of rye, where a minor difference was observed in replication dynamics of telomeric sequences. Telomeric regions of rye comprise large heterochromatic blocks (Appels *et al.*, 1978; Evtushenko *et al.*, 2016) and our results showed, that DNA loci closely connected to heterochromatic block (probably flanking regions) were replicated at late S phase, while telomeres located out of heterochromatin were replicated earlier, during middle S phase (Fig. 4B, D, E).

**Figure 4.**
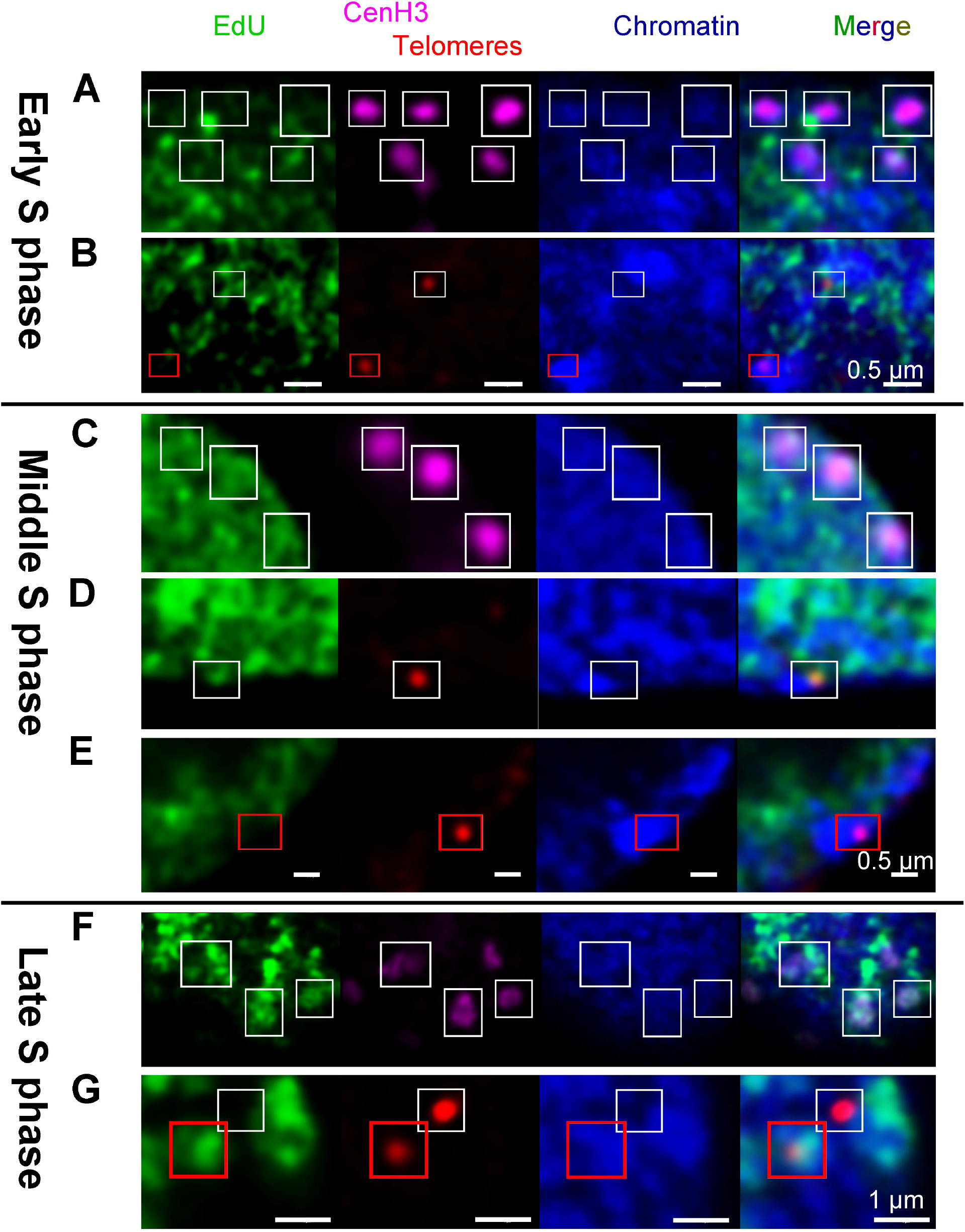
DNA replication in rye nuclei and replication timing of centromeric and telomeric sequences. Colocalization of signals specific to telomeres (red) and centromeres (pink) with EdU signals corresponding to replicated DNA (green) was used to describe the pattern of replication timing. In early S phase, centromeric signals (A) as well as telomeric signals which are connected with heterochromatin regions (B, red rectangle) did not colocalize with EdU. Telomeric sequences which were not localized in heterochromatin (B, white rectangle) colocalized with EdU signals, pointing to ongoing replication process. Colocalization of EdU and cetromeric signals is visible in middle and late S phase (C, D). Similarly, telomeric sequences connected with heterochromatin regions (red rectangle) colocalized with EdU in mid and late S-phase (E, G).

The difference in phasing centromere and telomere replication had an impact on the overall replication pattern of S phase nuclei as highlighted by EdU (Supplementary videos 1, 2, 3). The first replication signals in the nuclei were concentrated at telomeric regions, while the opposite pole of nucleus where centromeres were localized lacked EdU signals. In analogy, heterochromatin blocks and regions around centromeres replicated during late S phase, whereas the telomeres at the opposite pole of nuclei lacked the replication signals. A majority of nuclear DNA replicated during middle S phase, resulting in strong EdU signals spread across the whole nuclear volume (Supplementary Video 4, 5, 6). The images captured in rye are shown in Figure 4; supplementary Figures 2 – 7 show the images captured in the remaining species. The analysis of FISH signals by the Imaris software (Fig. 5) showed that co-localization between EdU and centromere fluorescence channels increased linearly from early S phase to middle and late S phase.

**Figure 5.**
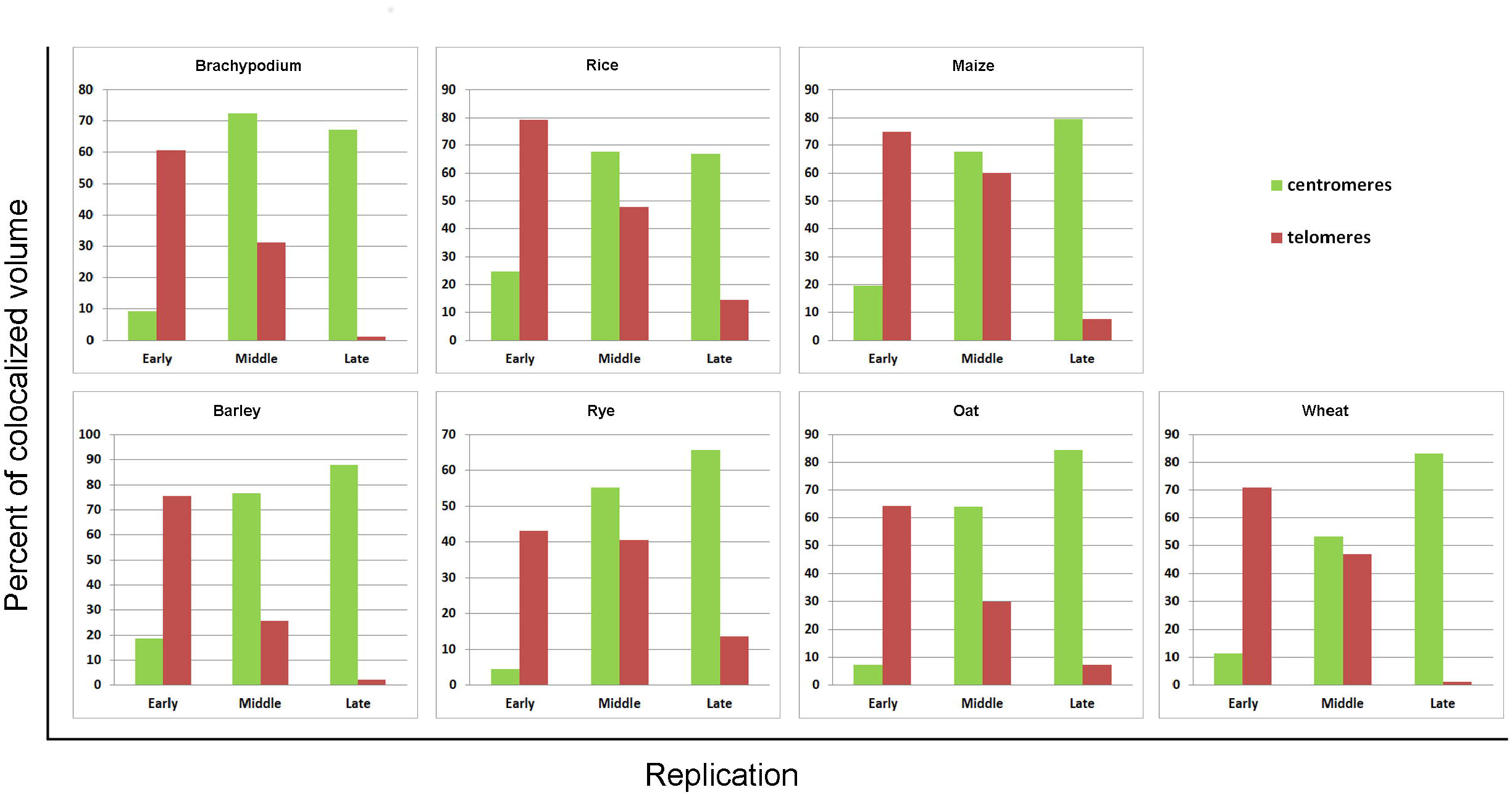
Replication timing of centromeric and telomeric sequences. Replication time was obtained after colocalization analysis and volume calculations of 3D models of microscopic images by Imaris 9.2 software. The columns represent percent of colocalized volume of signals from telomeric and centromeric regions and EdU signals.

#### Chromosome positioning in interphase nuclei

The localization of centromeres and telomeres during the course of S phase was used to infer chromosome positioning in interphase nuclei. In total, we analyzed 700 nuclei (100 nuclei for each species) with spherical shapes which are typical for the meristem cells of the root meristems. We confirmed regular Rabl configuration, when centromeres and telomeres localize at opposite nuclear poles, in large genomes of wheat, oat, rye, barley, as well as in *Brachypodium distachyon* with a small genome. On the other hand, chromosomes of rice with a small genome and maize with relatively large genome did not assume proper Rabl configuration (Fig. 6). The given interphase chromosome arrangement was stable throughout the interphase in all species (Fig. 7, Supplementary Fig. 8, Supplementary Video 1 – 12).

**Figure 6.**
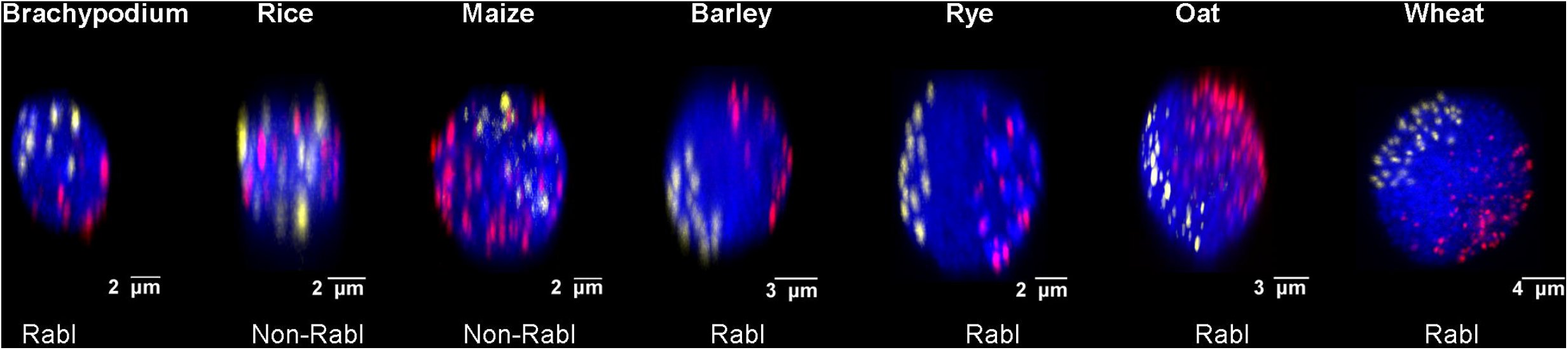
Chromosome positioning in G1 nuclei estimated based on the positions of telomeres and centromeres. Centromeres were labelled using CenH3 antibody (yellow), telomeres were visualized by FISH with an oligonucleotide probe (red color). Nuclear DNA was stained with DAPI (blue color).

**Figure 7.**
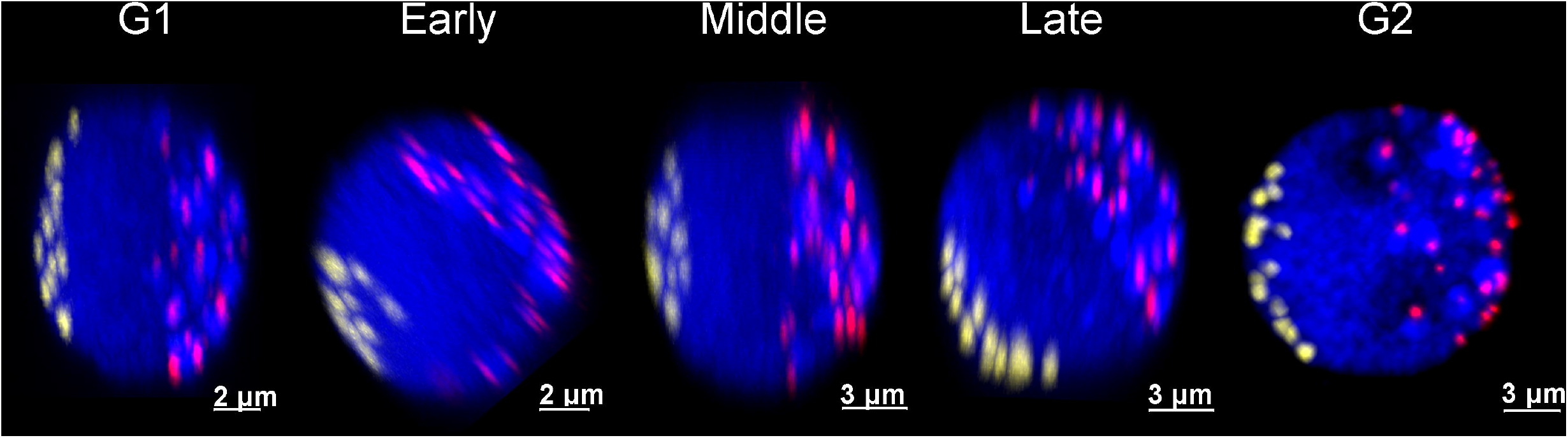
Orientation of chromosomes in interphase nuclei of rye. Centromeres were labelled using CenH3 immunostaining (yellow), telomeres were visualized by FISH with oligonucleotide probe (red color). Nuclear DNA was stained with DAPI (blue).

In a majority of species, the number of fluorescence signals from centromeres and telomeres corresponded to the number of mitotic chromosomes. The only exception was rice, where the Rabl configuration was not observed. Here, the telomeric and centromeric signals constituted large clustered signals randomly distributed in the nucleoplasm (Fig. 6, Supplementary Video 7, 8, 9). In maize, the centromeres clustered in one region of nuclei, but telomeres were randomly dispersed over the whole nucleoplasm (Fig. 6, Supplementary Video 10, 11, 12).

### Discussion

There is a growing interest to understand the principles of genome organization and its dynamics in three-dimensional space of interphase nuclei at various levels: from DNA fibers up to individual chromosomes and their domains. A range of studies focused on interphase chromosome positioning in plants (e.g. Pecinka *et al.*, 2004; Idziak *et al.*, 2015). However, except a study on maize (Bass *et al.*, 2015), chromosome positioning was not followed throughout the interphase, from G1 to G2 phase of the cell cycle. In order to fill these gaps, we analyzed spatiotemporal patterns of DNA replication and positioning of interphase chromosomes during cell cycle in root tip meristems of seven *Poaceae* species. They included important and evolutionary related crops differing in genome size and *Brachypodium distachyon*, a model wild species with a small genome.

A popular approach to study patterns of DNA replication in different parts of cell nuclei was microscopic detection of thymidine analogues incorporated into the newly synthesized DNA (Gilbert *et al.*, 2010; Bryant and Aves 2011; Bass *et al.*, 2014; Bass *et al.*, 2015). In mammals, early replication was observed in the interior regions of nuclei, while late replication occurred mostly at nuclear periphery (Li *et al.*, 2001; Pope and Gilbert 2013). In plants, differences in replication patterns at early, middle and late S phase were revealed by fluorescently labelled thymidine analogue 5-ethynyl-2’-deoxyuridine (EdU) (Hayashi *et al.*, 2013; Bass *et al.*, 2014; Bass *et al.*, 2015; Dvořáčková *et al.*, 2018). However, with a few exceptions, the patterns of nuclear DNA replication were analyzed only in plants with small and moderate genome sizes, such as Arabidopsis, rice and maize (Hayashi *et al.*, 2013; Bass *etl.*, 2014; Bass *et al.*, 2015; Dvořáčková *et al.*, 2018).

Earlier, Cortes *et al.* (1980) used thymidine analogue 5-bromo-2’-deoxyuridine (BrdU) in *Allium cepa* to observe a high coincidence between constitutive heterochromatin, including pericentromeric regions and late replicating DNA, which possesses a large genome. In barley, another species with a large genome, Jasencakova *et al.* (2001) found that DNA replication started at rDNA loci, continued at euchromatin and centromeric regions and was completed at pericentromeric heterochromatin. Our observations obtained after EdU labelling of newly replicated DNA agree with the results obtained in maize by Bass *et al.* (2015). Early S phase nuclei were characterized by localized weak signals, strong signals dispersed throughout the whole nuclei were observed in the nuclei at middle S phase and speckled signals concentrated in particular areas were observed at late DNA replication stages. As we studied seven species differing considerably in genome size, our results indicate that this pattern of DNA replication is general and does not depend on the amount of nuclear DNA.

We combined EdU labelling with the localization of telomere and centromere regions by FISH to provide a more detailed view on DNA replication kinetics. We observed opposite replication timing of telomeres and centromeres in all seven plant species, where the highest intensity of early replication was observed in gene-dense chromosome termini, while the highest intensity of late replicating DNA was typical for pericentromeric regions. In middle S phase, replication was almost evenly dispersed along the entire chromosomes and only slightly increased in the interstitial regions of chromosome arms. These findings confirmed the recent results made in Arabidopsis and maize obtained by Repli-seq analysis and corresponded to the gene density of their chromosome profiles (Wear *et al.*, 2017, Zynda *et al.*, 2017; Concia *et al.*, 2018). Kwasniewska *et al.* (2018) showed that terminal parts of barley chromosomes replicated in early S phase, whole chromosomes were covered with EdU signal at middle S phase and centromeric parts of chromosomes were replicated in late S phase. The authors reasoned that this chromosome replication profile implied presence of transcriptionally active genes in the terminal parts of chromosomes and inactive heterochromatin in the centromeric regions.

Two main types of arrangement were described for chromosomes in interphase nuclei of plants: Rabl configuration and Rosette-like structure (Rabl 1885; Francz *et al.*, 2002; Tiang *et al.*, 2012). The Rabl configuration, where centromeres and telomeres localize at opposite nuclear poles is considered typical for plants with large genomes. However, this does not seem to be a general rule and in some plants with relatively large genomes such maize and sorghum, Rabl configuration was not confirmed (Anamthawat-Jónsson and Heslop-Harrison 1990; Schubert and Shaw 2011; Tiang *et al.*, 2012). The Rosette structure, where the centromeres are located at the nuclear periphery whereas the telomeres congregate around nucleolus was described only in Arabidopsis (Armstrong *et al.*, 2001; Fransz *et al.*, 2002).

This work revealed a stable arrangement of centromeres and telomeres throughout the interphase. Based on the positions of telomeres and centromeres, we confirmed Rabl configuration in *Brachypodium distachyon* with a small genome, and in barley, rye, oat and wheat with large genomes. We also confirmed, that chromosomes in rice with a small genome and maize with a moderate genome did not assume Rabl configuration, confirming the results of Dong and Jiang (1998) and Santos and Shaw (2004) obtained by FISH on interphase nuclei. Some of the centromeric signals clustered at specific region of nucleoplasm, indicating a tendency to Rabl-like polarized organization, but telomeric signals were dispersed. One reason for the irregular distribution of rice and maize interphase chromosomes could be the presence of acrocentric chromosomes as hypothesized by Idziak *et al.*, (2015) who observed disrupted Rabl configuration in *Brachypodium stacei* and *B. hybridum* whose karyotypes comprise acrocentric chromosomes.

To conclude, our study indicates that spatiotemporal pattern of DNA replication timing during S phase in plants is conserved and does not depend on the amount of nuclear DNA. While the positioning of interphase chromosomes is stable throughout cell cycle, there seems to be more complex relation between interphase chromosome positioning and genome size. The observations by other authors that chromosome positioning may differ between tissues and even within tissue of the same plant indicates that interphase chromosome configuration is not a simple consequence of chromosome orientation in the preceding mitosis and that it is controlled by so far unknown factors.

## Acknowledgements

We are grateful to Dr. Kateřina Malínská for advice on confocal microscopy and we thank Ms. Zdeňka Dubská and Bc. Jitka Weiserová for excellent technical assistance. We acknowledge the core facility CELLIM of CEITEC, which has been supported by the Czech-BioImaging large RI project funded by MEYS CR, grant award LM2015062 for providing imaging facility. This work was supported by the Czech Science Foundation (grant award 17 - 14048S) and by the ERDF project “Plants as a tool for sustainable global development” (grant award CZ.02.1.01/0.0/0.0/16_019/0000827).

## Abbreviations

3D: three-dimensional
CenH3: centromere-specific variant of histone H3
BrdU: 5-bromo-2’-deoxyuridine
EdU: 5-ethynyl-2’-deoxyuridine
FISH: fluorescence *in situ* hybridization
PAA: polyacrylamide
rDNA: ribosomal deoxyribonucleic acid
ROI: region of interest
RT: room temperature

## Supplementary data

**Supplementary Figure S1**

Replication of 45S rDNA in oat. EdU signals (green) and 45S rDNA FISH signals (red) detected inside the nucleoli (white dashed lines) at early and middle S phase and in the periphery of nucleoli at middle and late S phase.

**Supplementary Figure S2**

Non-colocalization of CenH3 in early S phase in seven species. Immunofluorescent detection of CenH3 (pink) and EdU signals corresponding to replicating DNA (green) were visualized in *Brachypodium distachyon* (A), *Oryza sativa* (B) *Zea mays* (C), *Hordeum vulgare* (D), *Secale cereale* (E), *Avena sativa* (F) and *Triticum aestivum* (G). Non-colocalized signals are shown in white rectangles.

**Supplementary Figure S3**

Colocalization of CenH3 in middle S phase in seven species. Immunofluorescent detection of CenH3 (pink) and EdU signals corresponding to replicating DNA (green) were visualized in *Brachypodium distachyon* (A), *Oryza sativa* (B), *Zea mays* (C), *Hordeum vulgare* (D), *Secale cereale* (E), *Avena sativa* (F) and *Triticum aestivum* (G). colocalization is shown by white rectangles.

**Supplemetary Figure S4**

Colocalization of CenH3 in late S phase in seven selected species. Immunofluorescent detection of CenH3 (pink) and EdU signals corresponding to replicating DNA (green) were visualized in *Brachypodium distachyon* (A), *Oryza sativa* (B), *Zea mays* (C), *Hordeum vulgare* (D), *Secale cereale* (E), *Avena sativa* (F) and *Triticum aestivum* (G). colocalization is shown by white rectangle.

**Supplementary Figure S5**

Colocalization of telomeres in early S phase in seven selected species. FISH signals specific to telomeres (red) and with EdU signals corresponding to replicating DNA (green) were visualized in *Brachypodium distachyon* (A), *Oryza sativa* (B), *Zea mays* (C), *Hordeum vulgare* (D), *Secale cereale* (E), *Avena sativa* (F) and *Triticum aestivum* (G). colocalization is shown by white rectangle and is visible as yellow color in merged pictures.

**Supplementary Figure S6**

Colocalization of telomeres in middle S phase in seven selected species. FISH signals specific to telomeres (red) and with EdU signals corresponding to replicating DNA (green) were visualized in *Brachypodium distachyon* (A), *Oryza sativa* (B), *Zea mays* (C), *Hordeum vulgare* (D), *Secale cereale* (E), *Avena sativa* (F) and *Triticum aestivum* (G). colocalization is shown by white rectangle and is visible as yellow color in merged pictures.

**Supplementary Figure S7**

Non-colocalization of telomeres in late S phase in seven selected species. FISH signals specific to telomeres (red) and with EdU signals corresponding to replicating DNA (green) were visualized in *Brachypodium distachyon* (A), *Oryza sativa* (B), *Zea mays* (C), *Hordeum vulgare* (D), *Secale cereale* (E), *Avena sativa* (F) and *Triticum aestivum* (G). colocalization is shown by white rectangle.

**Supplementary Figure S8**

Models of 3D chromosome positioning in interphase nuclei of *Brachypodium distachyon* (A), *Oryza sativa* (B), *Zea mays* (C), *Hordeum vulgare* (D), *Secale cereale* (E), *Avena sativa* (F) and *Triticum aestivum* (G). The volume of nuclei (grey) was modeled based on the primary intensity of DAPI staining. Telomeres (red) were detected based on the intensity of fluorescently labelled telomeres and centromeres (yellow) were detected based on the primary intensity of CenH3.

**Supplementary Figure S9**

Maximum intensity projection of 3D nuclei from different stages of interphase in *Brachypodium distachyon* (A), *Oryza sativa* (B), *Zea mays* (C), *Hordeum vulgare* (D), *Secale cereale* (E), *Avena sativa* (F) and *Triticum aestivum* (G). All nuclei were counterstand with DAPI (blue) and early, middle and late phase shows replication pattern visualized by EdU (green).

**Supplementary Video 1**

Barley nucleus in early S phase. Centromeres were visualized using immunolabelling CenH3 (yellow), telomeres were visualized after FISH with an oligonucleotide probe (red) and replicating DNA was labelled with EdU (green). Nuclear DNA was stained with DAPI (blue).

**Supplementary Video 2**

Barley nucleus in middle S phase. Centromeres were visualized using immunolabelling CenH3 (yellow), telomeres were visualized after FISH with an oligonucleotide probe (red) and replicating DNA was labelled with EdU (green). Nuclear DNA was stained with DAPI (blue).

**Supplementary Video 3**

Barley nucleus in late S phase. Centromeres were visualized using immunolabelling CenH3 (yellow), telomeres were visualized after FISH with an oligonucleotide probe (red) and replicating DNA was labelled with EdU (green). Nuclear DNA was stained with DAPI (blue).

**Supplementary Video 4**

Rye nucleus in early S phase. Centromeres were visualized using immunolabelling CenH3 (yellow), telomeres were visualized after FISH with an oligonucleotide probe (red) and replicating DNA was labelled with EdU (green). Nuclear DNA was stained with DAPI (blue).

**Supplementary Video 5**

Rye nucleus in middle S phase. Centromeres were visualized using immunolabelling CenH3 (yellow), telomeres were visualized after FISH with an oligonucleotide probe (red) and replicating DNA was labelled with EdU (green). Nuclear DNA was stained with DAPI (blue).

**Supplementary Video 6**

Rye nucleus in late S phase. Centromeres were visualized using immunolabelling CenH3 (yellow), telomeres were visualized after FISH with an oligonucleotide probe (red) and replicating was labelled with EdU (green). Nuclear DNA was stained with DAPI (blue).

**Supplementary Video 7**

Rice nucleus in G1 phase. Centromeres were visualized using immunolabelling CenH3 (yellow) and telomeres were visualized after FISH with an oligonucleotide probe (red). Nuclear DNA was stained with DAPI (blue).

**Supplementary Video 8**

Rice nucleus in middle S phase. Centromeres were visualized using immunolabelling CenH3 (yellow), telomeres were visualized after FISH with an oligonucleotide probe (red) and replicating DNA was labelled with EdU (green). Nuclear DNA was stained with DAPI (blue).

**Supplementary Video 9**

Rice nucleus in G2 phase. Centromeres were visualized using immunolabelling CenH3 (yellow) and telomeres were visualized after FISH with an oligonucleotide probe (red). Nuclear DNA was stained with DAPI (blue).

**Supplementary Video 10**

Maize nucleus in G1 phase. Centromeres were visualized using immunolabelling CenH3 (yellow) and telomeres were visualized after FISH with an oligonucleotide probe (red). Nuclear DNA was stained with DAPI (blue).

**Supplementary Video 11**

Maize nucleus in middle S phase. Centromeres were visualized using immunolabelling CenH3 (yellow), telomeres were visualized after FISH with an oligonucleotide probe (red) and replicating DNA was labelled with EdU (green). Nuclear DNA was stained with DAPI (blue).

**Supplementary Video 12**

Maize nucleus in G2 phase. Centromeres were visualized using immunolabelling CenH3 (yellow) and telomeres were visualized after FISH with an oligonucleotide probe (red). Nuclear DNA was stained with DAPI (blue).

